## Supplementary Information for "Population genomics uncover loci for trait improvement in the indigenous African cereal tef (*Eragrostis tef*)"

**Supplementary Note 1: Additional Information on genotyping of the core collection with a minimal SNP set.**

We called the minimal SNPs on a VCF file derived from pooled, subsampled FASTQ files. We therefore checked that the individual constituents of the pools didn't disagree with the "consensus" calls for the 28 selected SNPs.

There were eight cases where the VCF file for the 220 individual accessions differed from the pool calls for the 28 SNPs. In six of these cases, one member of a group was called heterozygous where the group itself was called homozygous. However, in all cases these heterozygous calls were based on a total read depth < 10 and/or received a GQ score < 15. This implies that the calls were not very high confidence - indeed, the empirical filters derived for the 150 accession and 220 accession VCFs would have converted such calls to missing data.

Additionally, in each of these cases, the heterozygous calls for the group members trended towards the genotype assigned to their respective group. For example, DZ-01-348 received a heterozygous call for SNP 17, but the reads supporting this call were actually 1 read for the reference allele and 8 for the alternate allele. Given that the group that DZ-01-348 is part of (DZ-01-136 group) was called homozygous for the alternate allele, this suggests that a higher read depth might have led to the correct call for DZ-01-348 for this SNP.

Lastly, there were two cases where different group members had different types of homozygous calls for a SNP. However, these groups have missing data for those particular SNPs, so do not use them for discrimination from other accession groups and singlets. These discrepancies therefore do not change which of the 150 accession groups and singlets the group members would be assigned to.

For the new DZ-01 group,  DZ-01-383 is heterozygous for SNP 13. In the DZ-01-136 group, DZ-01-348 is heterozygous for SNP 1 and SNP 17. In the DZ-01-647 group, DZ-01-647 is heterozygous for SNP 7. In the DZ-01-1556_1 group, DZ-01-1556_2 is heterozygous for SNP 6. Also, SNP 6 has member accessions homozygous for both alleles, however, the DZ-01-1556_1 group has missing data here. In DZ-01-667 group, DZ-01-667 is heterozygous for SNP 7. In the DZ-01-1100 group, SNP 11 has member accessions homozygous for both alleles, however, the DZ-01-1100 group has missing data here.

### Supplementary Figures

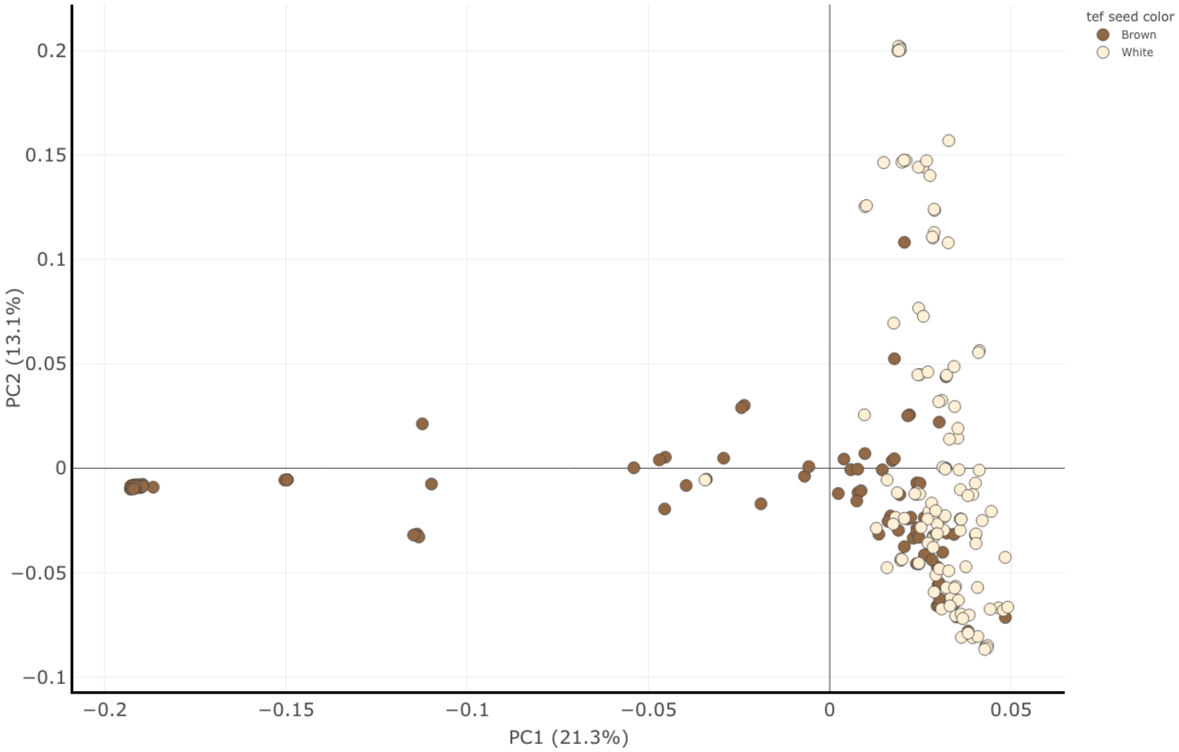


**Supplementary Figure 1 – Principal component analysis (PCA) based on ca. 40K SNPs for 230 tef genotypes**. A scatter plot of PC1 (explaining 21.3% of the variance) versus PC2 (explaining 13.1% of the variance). Binary colour codes represent seed coat colour of the individual genotypes

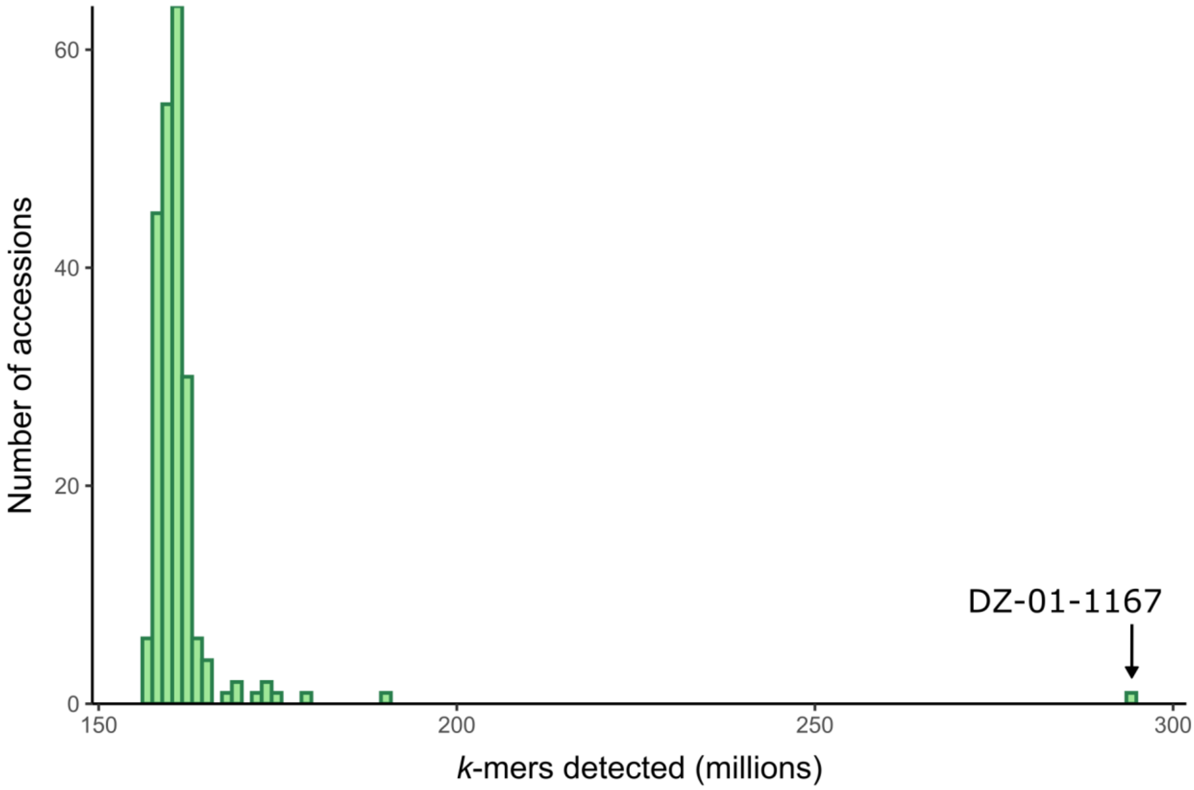


**Supplementary Figure 2 - Accession 'DZ-01-1167' contained a uniquely high number of distinct k-mers.** Histogram showing number of distinct k-mers per sequenced tef accession (before collapsing redundancy groups). Accession DZ-01-1167 displayed a significantly higher number of distinct k-mers than any other accession (84% higher than the mean of other accessions).

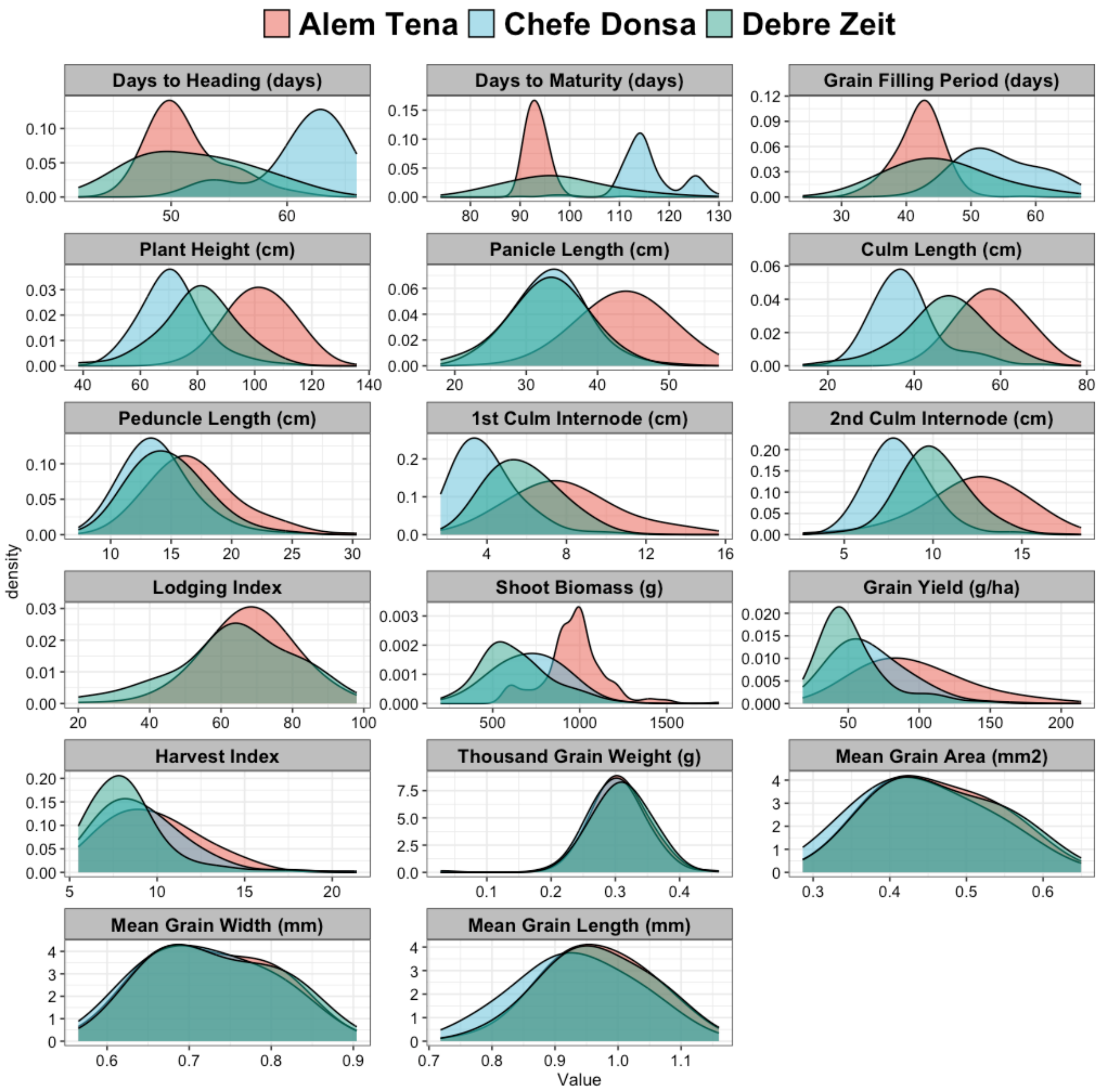


**Supplementary Figure 3 - Distribution of quantitative agronomic traits.** Density plots showing the raw distributions of quantitative agronomic traits for each location.

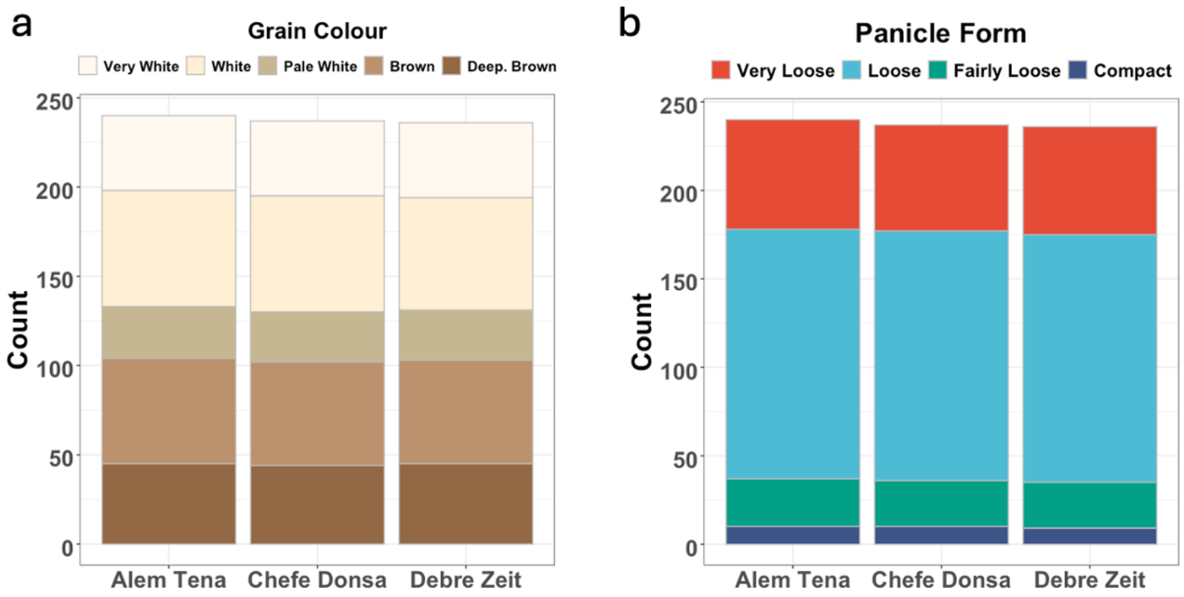


**Supplementary Figure 4 - Distribution of qualitative traits.** Stacked bar plots showing the distribution of grain colour (a) and panicle form (b) at each location.

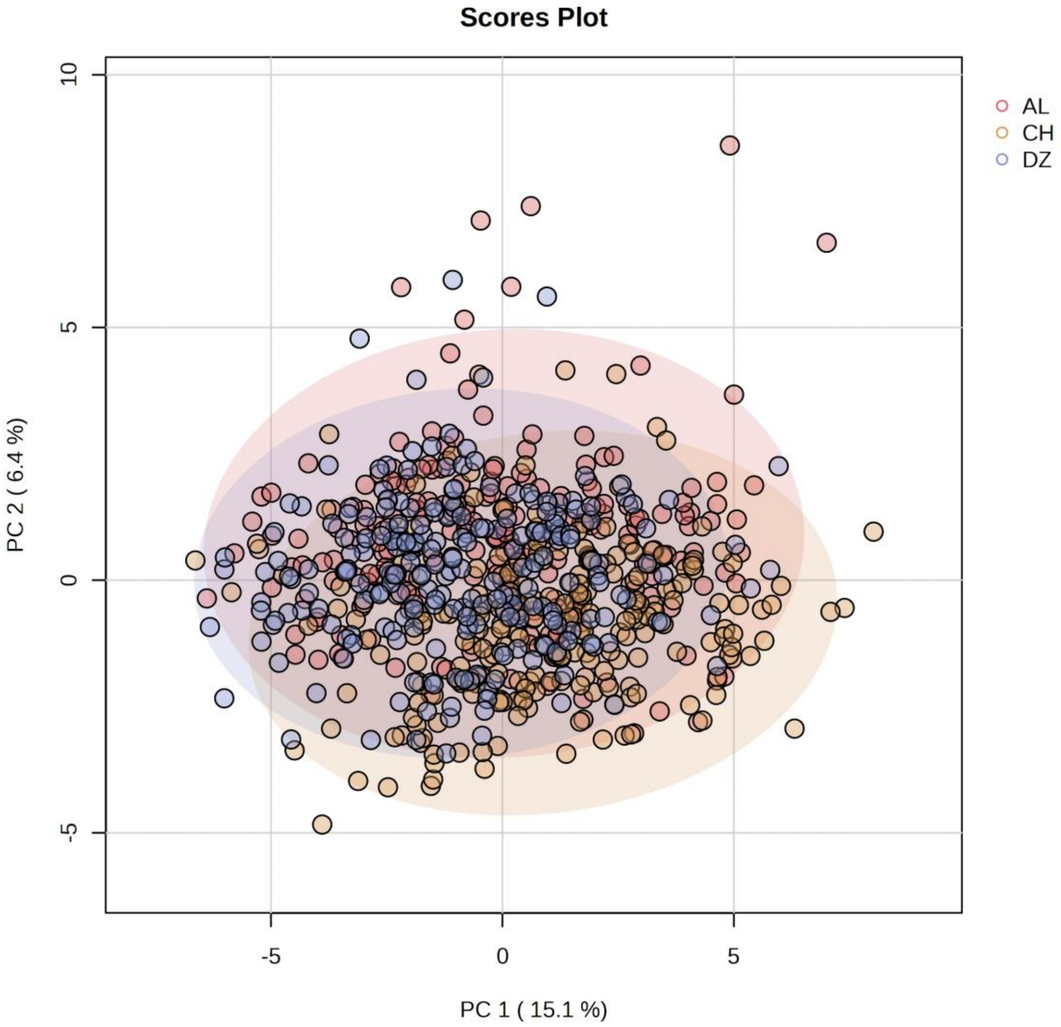


**Supplementary Figure 5 – Differentially accumulated metabolites between brown and white-grained accessions did not differ between locations.** Principal component analysis of trial plots from all three locations based on the 183 annotated metabolites that were differentially accumulated between brown and white-grained accessions. AL, CH and DZ represent Alem Tena, Chefe Donsa and Debre Zeit, respectively. Points from the three locations were not well-separated, indicating little effect of location on the grain accumulation of these metabolites. Ellipses represent 95% confidence intervals around each group.

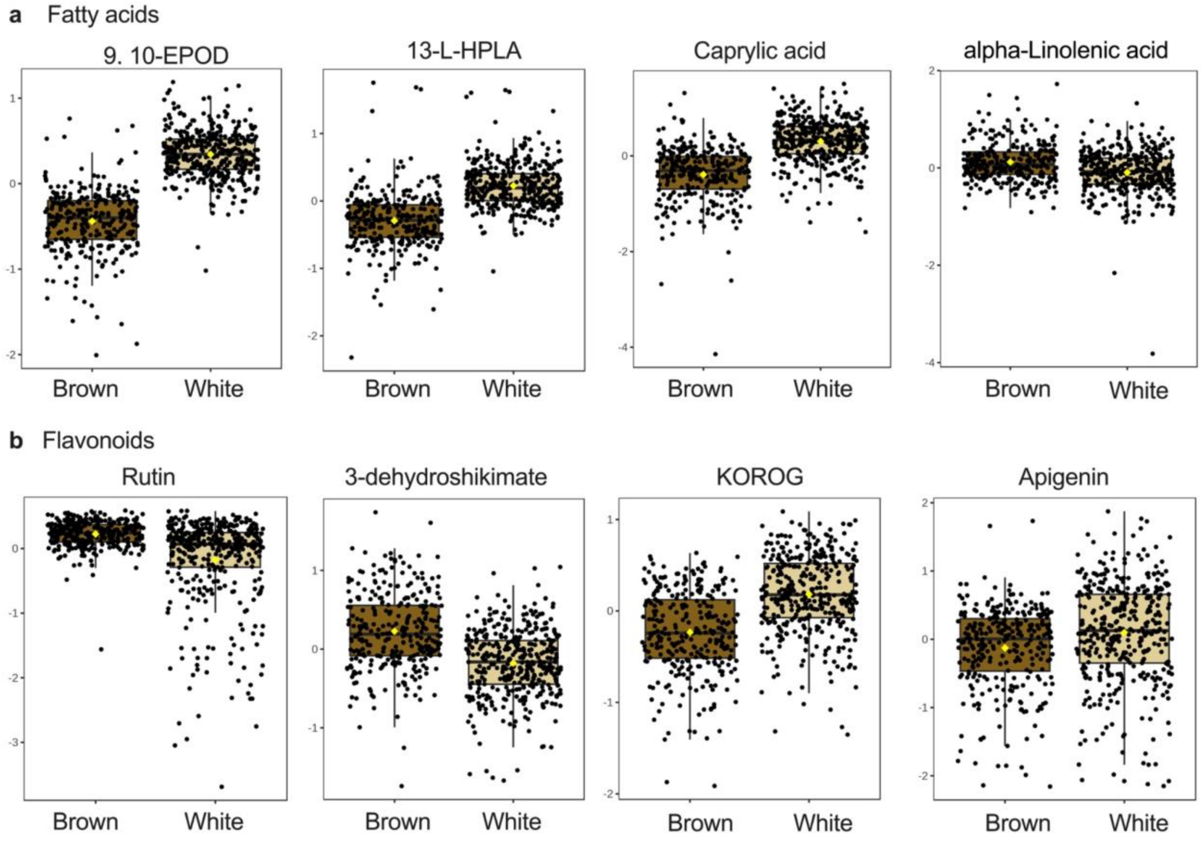


**Supplementary Figure 6 – Grain samples from brown and white-grained accessions differed in their accumulation of several fatty acids and flavonoids.** Box plots showing the accumulation of selected metabolites in brown and white-grained tef accessions. The metabolites in row **a** are fatty acids while those in row **b** are flavonoids. Metabolite concentrations are $log_2$ transformed.

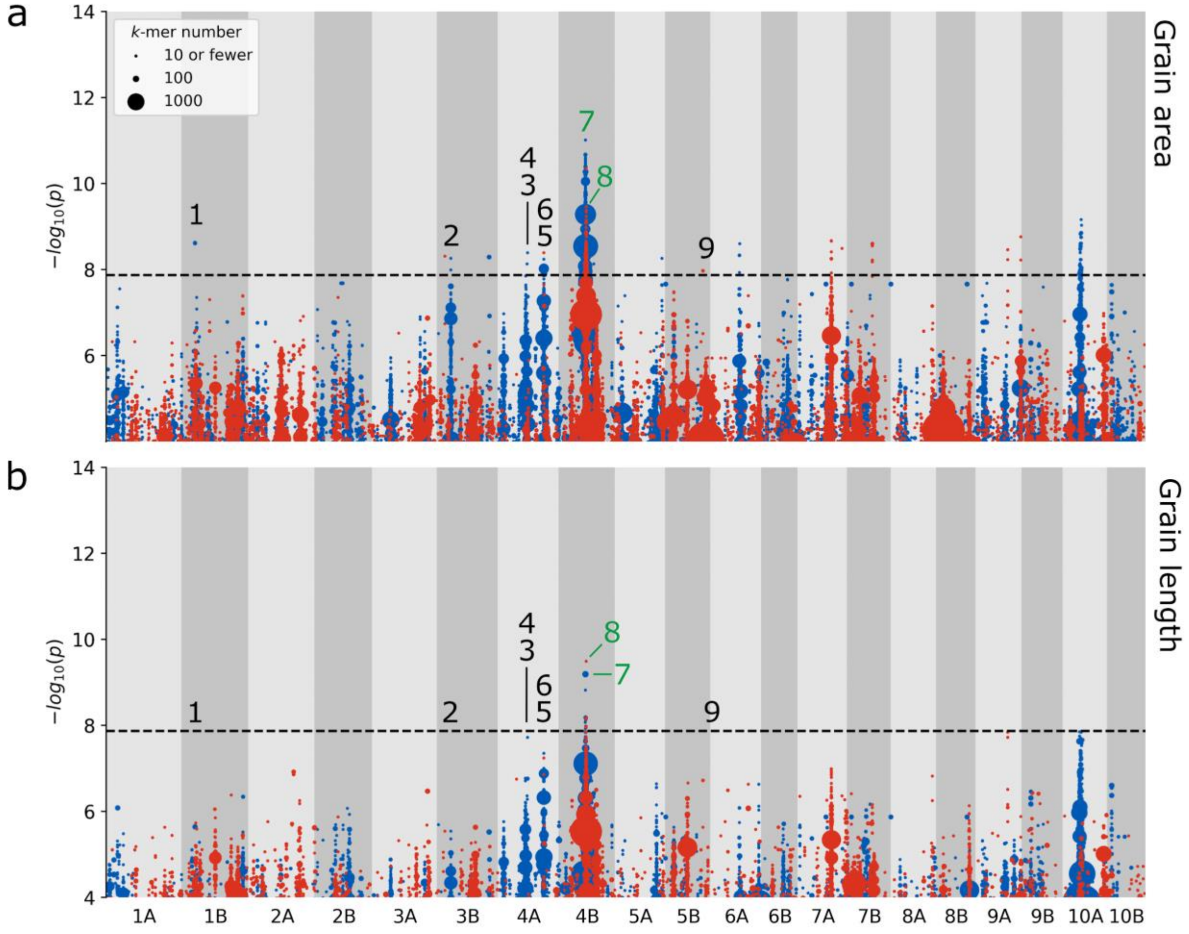


**Supplementary Figure 7 - k-mer-based GWAS identifies marker-trait associations for grain area, but not grain length.** Plots of *k*-mers associated with **a**, grain area and **b**, grain length. *k*-mers are grouped according to their association level and genomic coordinates (10 kb bins) and coloured according to the direction of association; red for association with lower trait values, and blue for association with higher trait values. Point size is proportional to the number of *k*-mers rounded upwards to the nearest 10. The nine highlighted regions from Figure 7 are labelled with black and green numbers, denoting whether the region is significant or not-significant for the plotted trait, respectively.

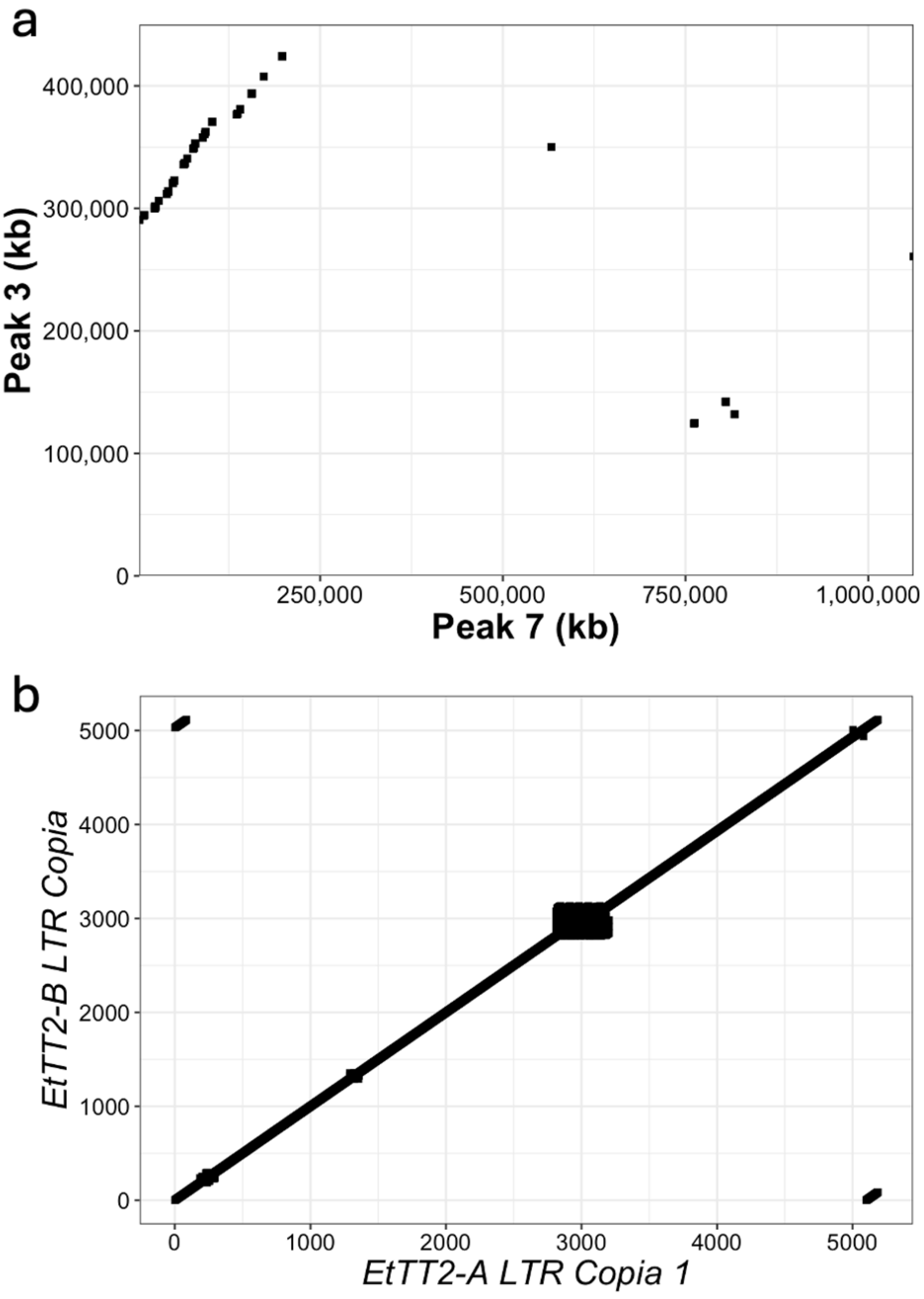


**Supplementary Figure 8 - Sequence alignments of chromosome 4 significant regions and LTR Copia insertions into *TT2* homoeologues. a**, Dotplot sequence alignment of two significant regions from chromosome 4A (peak 3) and 4B (peak 7) (window size = 100 bp, step size = 10 bp, match threshold = 90). **b**, Dotplot sequence alignment of the LTR Copia retrotransposons inserted in the A and B subgenome homoeologues of *TT2* (window size = 20 bp, step size = 5 bp, match threshold = 18).
