## Supplementary File 1 for "Population genomics uncover loci for trait improvement in the indigenous African cereal tef (*Eragrostis tef*)"

## > 4B:13922640-13939671

CGTCGGGATCCTTAGGGTTCATACAGTTTGCTGGTGTGGATCCTGGTCTAGGCTGATCCAGTGGAGGATCATCGTGGGTGTCGTGGTCTCGTGGACATAGAGATTAGGGGCAATGTGATTTTCTCAGATTTCGTGGTGCTTCTTTCAGGGTCTTTTATGAGTAGTTGGTGGTCACAAATCTAATCTCAGGAGATGATTGATGCTGTTGGTCGTAAACGATTTCTGGAATCTCGTTTTGGTGTGGAGTATTGGGGAATGGAACTAATTAAGTGATTTTTTGTCAATTTCGTCAATGATTGGTTTCTGTGATTGCGTCCCGCTTCAATCTTATGCTTAGTTGGTGTAGATTCTTCCCCAGCACACTGTAGTTGGTAATATAGTTGCGAAAAGAAGCTATGATTAGTACGTCAAACATGAATACATCACTCGATCCTGATGAAGTGTCATTGATATGTGTGCCCTAGACAGCAAAGAGGAAGGGGCCTTTTGTTGGTCTGAAGGCAATAAGGTGTTGAAGAAATTTCATCAAGGATCAAACCTTGTGTCGCCTGTGCCATAGCCTTTTTGTTGAAGTTGTTATGTAGGAGGACTTGGGTAGTAGCAGCTATGTTGGTTCATGTCTTGCTGATCAGGCAGTTCTGAATTTGTTGTGTTGCTGCAGGGGTAGGATGGTGGCTGCGGCCAATTTCTCCTGAGTGGTTGCAATGTGGTGTTGTTTGGTTAATTTTTCCCCTTTTTTCATTTTAAAATATGTCAGTCTTGCTTTAGTGGATAGAAGAATGAAGTCTTTGCTTCCATGTTTCAACTGGAAAATTTCAATCTGTTTTATTCGTTTGTATGATGTAAAACTGCTACATGACATGGAATGGCATAAGCGGTTACTTCTTTCATTGGTATGGGAGCCTAGCTTGCTTCGACGAATAGTTATTGTTGTTCCATGGAATCTTAATTTGCGGCAAGTTCTCATATCCTAATCTCTTCTTCTTTTTTTCCTTCTTTTTTTTCTGTGATGGTACTCCAATCCACCTAATGTTGTCAGAAATGCACTTTACAATTAGGTTGTATGTTTCTGCATAGTTTCGTTTTCATGGGGTAATATATTAGATTTCACATTCTATATAGAATATGTTTTTAGTCATGATATCAATTACAAATGCAAATATATCCATTGCATCCTTCAATTCTACTTGCATTAAAAAAATAAAAAGGCTATGTAGTCATCATGACTGAAAAGTATGTAGATATCCTTGTTTCATTAGTGACAGTGGTGATGTAGTGGAAAGATGTTAGTACTTGGCACAAAGCATGACCATCCATCCTTAGGTGTATTTTCCTCAATAAATATTTGCGTATCTGGATGTATTCAAATTTATGTTTGTTAGAAATGTCTGGACTTGATAACACTCATATTTATGTTCCTTGTCATCAGCAGTCCCCCGTCCTTTTAGTACTGCATAGCATATGGTGATTGTACAATTATGGCTATGAGATAGGTTTGTGAATAGCAACAGGATTTTTATCTTTGCCATGCATTTTGATATTTCTACTGTGCTTAGTATAGATATTGATAATGAATTTACTAAATGCGTTGGTGCATTTCTTGCTTTCTGTTCGAAACAATCAAGTGCAGTAAGATCATCACTAATATCCTCAATTTAGCTTTGTAAAATTCGGCTAATCATGTCCCTTTACCAAGCTGTGATTCTGTATTGATTGCTAATGTACGATTATTGGATGTTGCGCAGTGATGGAGAAGTATCAAAACAACGCTAGGTTTGCATCCTTTAGTGACGCTCCATTTGCTCTCCGCTGTAAGAAATCATGCGTGGATTTGTTATTCTTGTAAGTACTATGTTCTTTGTCATTCTGGTATAGTTTTCAGATGCATCCATTAGGAGATAGTGGAGACATAACTGGCATCTCTTCAAGTGCATTAGCCATTAGCTCATTTAGGATGTGATGTAATATGGAGCTTGAAGTGGCAATCTGCTCTGCTGGCACAGATCTGAACTGAGCTGTTGACCAGTGAAGCAATACATTTCCCATTTTTCATAGCTCAAGTAAATGTTGATTCCAGTCCTTTGTAACTCTAGGAAACATTGATAGTGGATACTGATCCTTTCCACTGTTATTGTGTGTGTGAAAGACATTATTATATAAAACGTTTGATTTTTGGAAACGTACGGAAGTGCTTGTCCATGCCTTGTGGCTTTAACCATTTCAGTCCTCTTTCTATTTTATCTGGATATTGGTATTATGTGTTGTACAGCTTCATGGGGAAACACCCATCTACTTTGAATCTGTGATTCTAAATTATCTTCTTTTTTTTTGTATGAGCTTTCGTCAATGCAGTGGGACCTCTACTATATCTGTTCTAAGTTTGTTGCCTTCTTTTCAGGTGATCTTGGTAGCTTGATCTCGAATCTTGAGTAAGCTAAGGGCTACGCATCCACAGGTGCTCTCGACCAAGGACATCTCCATTAGGAAGCCGGCCTCCATGAAGGTCAAGATCTTGCAGTGGCATGCGGTGGCTTCATGGACGTGGGATGCGCAAGATGAAACATGTGGCATATGCAGGATGGCATTTGACGGCTGCTGCACTGACTGTAAGTTTCCAGGTGATGATTGCCCGCTCATGTGGGGTGCGTGCAATCATGCTTACCATCTCCACTGCATACTGAAGTGGGTCAATTCGCAAACATCAACGCCCCTTTGCCCCATGTGCCGCACAGAATGGCAGTTTAAGGGCTAATGCACATTAGTGGAGTATCTGGAATGTATCAGCACGATGCAGCTTTCATTATTGGGAGAAAGCTAGGATGTATTGTTGGAATGGAATTTTAGCATGCTGTCACTTTATATTGTGCTAAAAGGCTACAATCCTGATTATCAGCATCTGCTTAAGTGAACTTGTGCCGACTTGAGGTTGTGGCCAAGCAAATGAAATGATAATATATCAGAGGCGTTAAGCTTTCTGACCTCATTGCTATTTTTTGTGGTGCTATTGTTTACTTTCTGAAGGATTTAAAGTGTCGACTTTGGGTTTTGTTCTGGTAAAGTAGTAACAAGAAGCATGAAGCTGTATATGGCTTTGCTTGCTTTTATCTCTGTTCTCTTCTTCAACAGAAATAGTGATCCATAGGTTATCAGGTGTCACTTTAAGCTCTGCACACAACTTCAGTAGGCATGTGCGTATTATTTAGTCGTATTCTGAAAAAGAAGTATCTAAATTTTGTTTAAAGAAGGAACGAACGGGACATGCAAATTATGCTATCGTGGTAAATTATATCTATTCCTTCCAAAACTTTGTCCCTTGCAATTTGTTTCTTCCGTGCCAATGTAGTAATGGCAATATGCCAGCAAATTATTCCTTATTAGCTTATGCAGAAGCATGGATTGACTGTTAGGGCCGGTAGAGCTTTTCTAACGGCTTCACATAATTGTCAACCAAAGGGGTGCTTTTTCAGATGCTTCTTCTTAAAAATAGGAAGAAGCTTTATCTAAAAATAGGTTGAACCATGAAGCCACAAAATGGTAGCTTCACCGGCTTTCTTTCACAACCGCGCTCTCTACTTTACATATTTGTCCCTAAAAATAAGAAAAAGCTGCTGGAAAAGCCGTTTTGCCAAATGTTTTGGGCACAGCCGTAGACACAGCGAGCACAACCGTTTTTCTAAAGAAGCCACAGCCACAGCCGAAGCCGGCACAGCCATAGCCCTACCAAAGAGGCCCTTAGTGCTTAGTGCATTGAGTTACTTTCCCCATTAATAGGGAGGCTTTCTGTATGTCTCATAGTTTGGTTCATATGAAAATTGGATGTATAGCAGTTACACGTTCATAGACTTGTTTTGTTACTTGTCCAAAAATCACTCAAAATGAATGAAAATTGTTGCACTTGGTGTTCCCCCCCCCCCCCCCCCCCCCCCATCTGATTAAGATATATGGAGCCACCGCAAATATTTGTTTGAGATCTTGTTTTGCATGCAATTGTAAGTTCGATTACCATTAAGGATATATTGACAGTTAAAAAATTGTTAACCTTTGTTTGTTGGACGGACCCCATGCAGCCTTGGGCTTGTGCTAGGTTCCGTTGGCAGCCAAACTAAGCCAAATCGCCGGCGAGAAGTTTGGCCAATTGCACTGGGCACCAAATTTTTGGCAAGGTTGAGATCATGAGATGGGCAGTAAACCAAACACGCACTTGGATCTGGGGGTAACGTAGGTGTGCTAACAGTACATTCTAACCCGAGTGACAGCGCAACATGCAACATTTGCAAGCAACCAACATTTTTCTTCCTAGCCAAACGATTGAACACACCGGGTATAAACCAAATTAACTTGTTACCTCCTGTAAAAACAAAGCCCGTCTAGCTCAGTCGGTAGAGCGCAAGGCTCTTAACCTTGTGGTCGTGGGTTCGAGCCCCACGGTGGGCGTTATCTGTAGCACTCTTTTTTTGAACCCTATCTTTTCTTATATGATTACTTCAAAAATATACAGTGGAATTCTGGAATGTCACGTGCTTTTGTTAAATTTGTCTCTTCGTATGATAGTGCAAAATACGCTGGAATTCTAGAATGTCGCATCCTTTCGACTTGTCCGATTTGCTTCTTTATTCAGATCATGCTGGTCCTCCTCGATGAGCTTAACTGGCATCATATGTAGCCAGCCATATCTAATCAGCGTACAGTAAGATGATGATGAGCTTTGCCCTTAAATCCTCACTAATTCCCACCACGTCGCGGTACAACCTATAGCAACATCAAAACCATGCAAACTCCACGTCATCTCCTGCCGGCTCCAGCAGCAGCGCCTCCATGTCACCCTCGCCGCCGACGAACTGGCCGGCCGGCCCCACCACCCCGTCGCCGGCCTCCCCGTGCCACGGGCTCAAGAAAGCCAGCTCGTCCATGTCGAAGTCAAGGTCGATGGACACGTCATCCTCCGGCAAATGCTCCTGTAGCACCTCCGCCGCCGCCGGCGCCTCCGGTGGATGGCCGTGGCTCTCGGCCGGCGAGGCGGCGGCGCGCGGCAGCCTGGCGGGGCACCGCATCGCCTTGGTCCGGACCGGTGGGCTCGTGTTGCCGCCGCCGTCCGGCGACGGAGGCGCGGCGTCGCTGCTGGTTCTCGCCGGCTCCGACGACGGCCGGCTACGCACGACGACGGGCGACGCTGCGCTGGCCTGATGACACTGTTTCGAGTGGCCGCCCACCTCGCCGCGCACCTTCTTGCCCAGCGTCGTGTTCCAGTAGTTCTTGATCTCATTGTCTGTTCGCCCCGGCAGCCGCCCGGCGATCAGGGACCACCTGTCCGGCGACGACGACAAAATATGCAAACACGTACGAAATCGTTAATTAGTCACTGCTGGATTGCAGTTTCAGAGACATAGCATTAGATGTGTCAATCATCGATGGTGCCTGCCAGCCTGCTGTTACTCGATCTGTCCCGTTGATCACATCAAGTTAGGAAACTAATCACATCAAGTTATTAGCAAGTAGATACGAAATACTTGCATTGGCAGCTAGCACAGTTAAAGTTTCCCATTTCTGAAACGACAACGCTTTTGCGGTCAGTGAAATTTTTTGGAATTTTCAGGGTGAAGGAAACGCGAGTTAGGAAGTGGTTACGCGCGCCAGAGGCCTTTCCTGAGTGCGAGATATTCTCCTGTTGCTGTATATTTAGGGAGGGAGGGACAGGGCTACTAGCTGTCATCATCAACTTTGCTGACCTAATAACTGGTTCATCTTCAGAGGGTAGGACTAGGAGACACTGCCCTGCCAACGAATACGTGATCTTCTAGAGCGCGAGAGCTAAACAGGAAAAAAGAGGTGCTGACTGATATGGACAAAGTGGCAGTAGTGGTTATGATAACTTGAGGCAATTGGCCGTTGTGCTTGCCTGCTGGCCTACTTTGTCTGCTCAGATCTCTACCACAGCAGGACAAGGAAAAGTTTCCAGCAGAAAGAGTTTAACATGCAAGCTAATGTGTTGTATGCATGAATAACGCAATTACCCTTCTGTCTTCTATACCATACAAACTAACATTTCTTTGAACAATAACAAACCAAATTTTAGTATCTTTAGGATTTAATTTGCGAAATGCAAATCGATGGAAATGATCGAAATATGGATAACCACATACCTGTTGCCGAGGAGCATGTGCAGCCTGATGATGAGCTCCTCTTCGTCGTCGGAGATGTTGCCCCGCTTGATGCCCGGCCTAAGGTAGTTGAGCCACCGCAGCCGGCAGCTCTTCCCGCAGCGCTTCAGCCCTGCAACACGTCAATGTCGTCAGTGAGGCCGTTGCGCATTCCCAACAGTCAGCGAATTACTGTACTGAATTTGTCATGTCACATTGCATTTTGTCCTTGCATCGCCCTACTTGTAACCGATCCAAGCGTACTTGCTCTCCGTTCCAGTTACAGAGGTTTATGTAGACTCGTATGGTAATCTAGTCAGCATTGATTATGCCTTTGATTAACAGTTTGTTTAGCGAATACGGTGAAAAAAAAAAGACCTGTTTTCATTTCATTTTTATCCTAATATCTGATCTTAATGTAACTTAGCATGCATGAACTACAAGACGTGACAAATCCATACATAGATGACGATAAAAATCAGAAATGACTAAAAAGAGAAGATAAGAAACAGGGGACAGGAGAAGAATTTCAGGATCAGACCAACTTCCCGCATTGTGAATAAATTGATGTTGATGACGAGAAATCAGCAAGAGAAAGGGAGGAGGTGTCACTGACCAGCTCTTTTGGGGAGGCATCCCCACTTGCCCTCTCCGTGCTTCC***TGTTG*GAGTGTGTTGGCCCATATAGAGGCCCATGTATGTTGGATATATATATACCCACCCATCTAGGGTTGGAGGAATCAAGTCGTCCAGTACCCTAACTC*TAACA***TGGTATCAGAGCCTCTCTCTCTCCCTACCCAACCCTAGCCGCCGCCGCCGCCATGGCCGGCTGGCCGCCGTCGGGATCACCGCCCCCCGTCGTTGCCGCCGCAGGTGCGTGCGCTGCCGCAGGCCACCCGCCGCCGCCCCCCGCCGTCGCCGCGGGTGCGGGCGCTGCCACGGGCATGGACCTCGCGCCGCCGCCCCCCGCCGCCGCTGCAGGTGACCCACCAGCCGATCCTCGTCTCCGACTCGATGCCAGCGATCCGCTGGTGGCCCAGCTCCACCTCCAGGCAGTCGGTGTCCAGAACGTCCGCGCTATGGTCTCCATCGTCCTCGACGTGTCGTCCACCAACTACGGCCGTTGGCGCGATCAGATCCTCCTCGCCCTTCGCCGCTACGCGCTCGACGACCACGTCCTCACCGACACTCCTGCTCACGCGCGCGGTGGTTCGTGGATCTACCTCGACAGCGTCGTCGTGTCCTGGATCTTCGGGACCCTCTCCCCAGATCTTCAGGAACTCATCCGGGCTTCTGAGGACACCGCACGCCAGGCCTGGCGGGCTCTCGAGAACCAGTTCCTCGACAACGCCCAGGCCCGCGTCCTACACCTCGACACCCAGCTCCGCACCTTCTCCCAGGGCGACCTTTCTGTGACCGACTACTGCAGGAAGATGAAGGGCATGGCGGATGGGCTTCGGGACCTCGGTTCTCCTGTGGACGAGCAGCTTCTGGTCATGTACCTATTGAGGGGTCTCAACAGCAACTATGACCTCCTCAAGACCTGGCTACCTCGACAGCGCCCCTTCCCCTCCTTCCAGGACGTGAAGAATGAGCTCCGCTTCGACGAGATCACCCGGGGCCTTGCACCAGGACCTGGATCCTCCACCACCGCCCTCGTTGCTGCCCCGCCCACCACTCCACGGGTTGCCGCTCCTCCCCCTGGGCCGAGCGGGGACGGGGGTGACCATCGTCGTCGCCGGAGACGTGGGGGCGGCCGCGGGACCTCTACACCGACTCCTACTCCGGCGGGTCCGTCCTGGCCATCCTTCACCAACCCATGGTCGGGGCGCATCTCCATGTGGCCGTTCCAGCCCGCCGGAGCCGGGGCTCGTCCTCCGCACCGGCCCGCTGCTATGCTCGCCGGCCCTGCTCCCCCGCTGCCCTGGGCTCCTCCCGCTCCGGTCGGCCAACCTCCGACCTGGCCGGGGGGGTGGACTCGTCCTGCTCCGACTTGGCCGGGGGGGTGGCCTCCTCCTGCTCCGATTTGGCCGGGGGGGCAGGACCCGGCGGCGTTGGCCCAGTCTCTCAGCACGATGCCGCTGACCCCACCAGCCGACTGGATCGCGGACTCGGGAGCCTCGTACCACACCACGCCCGATGCCGGTATACTTTCTTCTGTCCGACCTCCCCACCCTTCCTGTCCCTCTGCCATTATGGTGGGCAATGGTGCCTGCCTTCCTGTCACCTCTGTTGGCACTGTGCCGGGTTCTTTTCCCCTTTCTGATGTTCTTGTTGCTCCCCAGATGGTTCACAACCTCCTTTCCATTCGCAAATTTACTACCGACAATTCTTGTTCTGTTGAGTTTGATCCTTGTGGTGTTTCTGTGAAGGATTTGGCTTCCCAGCGTCCACTACTTCGATGTGACAGCACGGGACCCCTTTACACGATCCGCCTTCCCGCTTCCGCTCCTTCATCGCCATCTTCTTCACCATTGGCCTTTGCTGCTACATCTTCTTCTACCACCTGGCATCGCCGGCTTGGACATCCCGGCCGTGACGTTTTGGCCCGACTCTGCCGCAGTGTGGATCTCTCTTGTACTCGGGCAGATGATGAGCACCTCTGTCATGCGTGCCAATTAGGTCGTCATGTTAGACTTCCTTTTCCTACTTCCCAGTCGCATGCGACTCATGTTTTTGATCTTATTCACTGTGATCTGTGGACTTCACCTGTAGTCAGTATTTCTGGGTATAAGTACTACTTGGTGATTGTTGATGACTTCTCTCACTATTCTTGGACTTTTCCTTTACGCGCGAAGTCTGACACTTTTCCCACCCTCCTCAACTTCTTTGCTTGGGTCTCCACTCAGTTTGGCCTCACCGTTAAAGCTGTTCAGTGTGACAACGGCCGTGAGTTCGACAACCACACCTCCCGAGCCTTCTTCCTCTCCCGCGGCGTTCACTTGCGCATGTCTTGTCCGTACACATCTCCTCAGAACGGCAAGGCAGAGCGCATGATCCGCACTACGAATGATATTGTTCGTACCCTACTTGTACAGGCCTCCCTCCCGGCCCGCTTCTGGGCCGAGAGCCTACACACCGCCACCTATCTCCTCAACCGTCTCCCTTCCACTGCCTCCCCTGCTCCCACTCCCCACCACGCCCTTTTCGGCTCCCCTCCTAGATATGACCACCTCCGTGTCTTCGGGTGCGCCTGCTACCCCAACACCTCTGCCACTGCTCCTCACAAGTTGGCGCCCCGTTCGACCCGGTGTGTATTTCTCGGGTACTCTCCGGAACACAAGGGGTATCGCTGCTATGACCTTACATCTCGTCGGGTCCTAATCTCTCGGCACGTCGTGTTTGATGAAACTGACTTTCCCTACTCCACCTCCACCACTCCCTCTCTGGATCCCGAGATCGGGTCTGTGTTTCCGACTGACCCGGTGGCCCAGTCACCTGTCTCTGTGTATCCTTTACCTACAGGTCTTCAGGGCACCCCGACCCCGAGTCGGCCGCGCTCGCGATCCCCGCCTCGCCCGACCCCGAGTCGGCCAGGTCGGGCGGACCCCACCCCGAGTCGGCCGCCCTCGCGATCCCCGCCTCGCCCGACCCCGAGTCGGCCAGGTCGGGCGGACCCCACCCCGAGTCGGCCGCGCTCGCGATCCCCGCCTCGCCCGACCCCGAGTCTGCCGGGCCGGGCGGACCCCACCCCGAGTCGGCCGTGCTCGCGATCTCCGCCTCGCCCGACTCCGCCTCGCCCGACCCCGAGTCGGGCGGGCTCCCGACCGCCCCCGAGTCGACCGGGCTCGCGATCCCCGCCTCGTCCGACCCCGACACCTTCTCCCGCGCGCTACGCCCGGCCTCCGCTCGTCTACCAGCGCCGGGCGGCACCAACTCCACCGGTGCCTCCGCCCCCACAGCCGACTGCGGCCCCGTCTCGGTCCCTCCCCCGCCTCAACCCGGTGGTGTACCACCCGCCTGTCATCCATCGGGATCCCGGACACACTCATCCCATGGTGACCCGGCGGGTGGCGGGGGTTCTTCGACCTAAGGTCCTCACTGCTACTGCGGACGAGCCGGCGGTCTCGCCGATTCCTACCTCTGTCCGTGGCGCCCTGGCTGATCCTCACTGGCGCCGGGCGATGGAGGAGGAGTATGCGGCCTTACTTGCCAACCAGACGTGGGACCTCGTTCCTCGTCCACCTGGTGGCAATGTGGTCACAGGGAAGTGGATCTGGACTCACAAGCGTCGTGCTGATGGCACACTAGATCGGTACAAGGCTCGCTGGGTTCTCAGGGGATTCACCCAGCGGCCTGGAGTTGATTATGATGAGACCTTCAGTCCCGTGGTGAAGCCGGCTACAGTCAGGTCAGTTCTGTCGCTGGCTCTCTCTCGCTCTTGGCCTGTGCATCAGCTGGATGTGAAGAATGCTTTCCTGCACGGCACTCTCTCTGAGACAGTTTACTGCAGTCAGCCAGCGGGGTTTGTGGATCCTGGTCGTCCTGACATGGTCTGCCGACTCAACAAGTCCCTCTATGGTCTGAAGCAGGCACCTCGGGCTTGGTACTCTCGGTTTGCCTCATTCCTGGTCTCTTTGGGTTTCACTGAGTGCAAGTCGGATACCTCGCTGTTCGTCTACGGCCATGGCGCTGACACTGCATATCTACTGCTGTATGTTGATGACATCATCCTCACAGCGTCCAGCCCGGCCTTGCTCCAGCGGATCATTCAGGCCTTGCAGCGCGAGTTTGCTATGAAGGATCTGGGTCTCCTTCATCACTTCCTGGGGGTCACTGTTGAGCACCGTCCTTCTGGCCTGTTCCTTCATCAGCGGCAGTATGCTCTTGATATTCTGGAGCGTGCCGGGATGTCTGACTGCAAGCCGAGTTCCACCCCCGTTGACACTCAGGCCAAGCTATCTGCTGATTTGGGCGCTCCGGTTGCTGACCCGACTGCCTACAGGAGCCTCGCTGGGGCTTTGCAGTATCTCACCTTCACCAGGCCTGACATCGCTTATGCCATCCAGCAGGTCTGTCTTCACATGCACGACCCCCGCGAGTCCCACCTGGCTGCTCTGAAGCGCCTCCTCCGCTATGTTCGTGGCACAGTTGACTATGGCCTGACTCTTCACCGGTCTCCTTCTGCTGAGCTGGTTGTCTACACGGATGCCGACTGGGCTGGCTGTCCGGACACTCGGCGTTCCACCTCGGGCTACGCAGTCTTCCTCGGCGGCAACCTTGTCTCGTGGTCGTCCAAGCGACAGCCCGTGGTCTCCCGCTCCAGTGCCGAGGCGGAGTACCGGGCTGTGGCCAACGGCGTCGCTGAGGCGACCTGGCTTCGACAGCTCTTGGCGGAGCTCCACACTCCACTCGCCAAGGCTACACTTGTCTACTGCGACAACGTCAGCGCGGTGTACCTCTCCACCAACCCGGTTCAGCACCAGCGGACGAAGCATGTCGAGATCGATCTCCACTTCGTTCGGGACTTCGTCGCCATTGGGCAAGTGCGGGTCCTCCACGTTCCCACTACCTCCCAGTTTGCGGACATCTTCACCAAGGGTCTCCCCTCCTCGATCTTCTCGGAGTTTCGATCCAGTCTCAACATATCCGGTGGCTAGTTGTGACTGCGGGGGGGGGGGGGGGTGTGAGTTGTTCTTTCTTGTGTCCAGTCCTGAACACCGCTGCGACGGTTAGTTCAGACTGCGGGGGGG***TGTTG*GAGTGTGTTGGCCCATATAGAGGCCCATGTATGTTGGATATATATATACCCACCCATCTAGGGTTGGAGGAATCAAGTCGTCCAGTACCCTAACTC*TAACA***CTTCCTGATGTAGGAGACGAGGATGTCATCCTCCATTGCTGTCCAAGCTCCCCTATTCAGCCCCTCCTTGGGGCAGCATGGTTTCCTCCCCATCTCTCTCTCTCTCTCTCTCTCTCTCTCTCTCTCTCTCGGTTAGATGGAGGGTTATGCTCTGTGATTTCCACTGAGGTCCCTATTAAACAAGGTGCAGAGCGCTCCGTATTCCATGCCCATTTTTTTTTAAAAAAAAATATTTGTGATATAGCCCCTCGAATGGACATTTATGCCCTACGGGCACTCTGTTTTCGCCCACACAGCTACTTCTGATCAGATGCTCGTTATGAAGGCAGTTTCTAATCGGACATCAGGAAGCTTTCGCTTGTTTATGCACATATAGGGAATGCTTAAAATTTTGACCAGCATAAAATTTCAGAAAATGATTTACAAAACACCGAATTACTTAAGACCATATCGAATACGACAAGTGCATGTTCTAGATGCAGATGCCAATGTGGCACCTGCAACAGGCTTGAGAAGTACCGAGAGGGAAAACATATTGACATACAGTTTTTGTCAACTCTGATTCTTCTTCGTTTATCAAGGTTCAGTAGCCATGGTAATGAAAAAAAAATCTGATCTACAACCATGGTGTAGGAAACTACATCCTATCTGGGAATATACTCCTACTAATTCACAAATAGTCAACACTTACCTATTAACATAAATCTAAAAGGTGGATAATAGCACAGAATTTATTTGCAAATGCAACAAGCATAACCTTAGCTAAAATCCTTAAGCCTTCTCTTCAAAACATCCATGGACCTCTTGCGAGCACTCTGCTTAGCGTTACGGTTCGAAAGGCTCCCAAACAGCCCGTCAGCAGCGTCCTTGTACTTGTCACAAATGATGGAAAGCTTTTTGTAGTCCAGTGATCCGTAGTTCCTCTGTCTTATCACCGGATTCATGAAGTTCTCTGGTTGGCTACTGAGCAAGAGGTTGACTAGCTGCCTTATTTCAACCAAACAATCTCTGAAAGTTGTTTCCTTTCCCGAAACCGACAACCCCATGCTTTCGAATTTCTCATCTGCAAATGCCTCCAGCATTTTCAAGTCCATATCAAGACCTAGAACTGCATTAACAGTGAACCTTTTCACACCTTCGTTAAGAAGTGTTGTCATTATGGAGTCTGAGATATGGCTCATTATACCAGAAACCACTTTGTACAGAGCTTCCAAGGGCAAAATTTCCTGTGCTGTTGATACTTGAGTTTCGAGATATATCAGGACCTCATTCATATAGTCGTTTGAAATGTCTGGGGCTTCCTCTGCTATCCAGTTGACGTTTTCCAAGAGTATCATAAACTCATCGACCTTGAAATTGGCCAGGTTTATCAGAGCATTGTATACTGCATTCTGCGAGGCTTTCAAGACAGCTCGAGCAGTTAAACTAGAATGAGATCTCTCCGCAATGCGTTTTGGTATGCCACACAGATGAGCAGCATGCAAGAGGAACATGCCACATGCTTGCTCAAGAACAGAAATGTTACCAGCGAGCTGCATCATCTGTGACATGGCCAATGAGCGAGCATAGATCATGTTCAGAAGACTGTCATTTAAGACCTCAATCAACAGCCTGTCTAGGTATGCTTTCACAACATCATACATGTTCATGAAGCCACCATATGACAAGTAGCTGACGGAATCTTCAATGAAAGACCGAACGATGCGACAAATTTCTGGAACAGAAGAGGAGAACTGCGCCACATATGGGAAATCAGGGGCTGCATCATCAGGCTCCAAACAAAATGCTGTCACATTCATGTTATATTCGTATTCCTTCTTTATAACCATCTGCTCATATGAGTCATTAATAAGGATGTCATCTATCTGCTTTCGGCATTCAAGAAGAAGCAACTGGTGGTACTTGTCCCTGCTTTTCTCAAGAACTTCAATCAACTGTGTAATTTGATAACCATATTTCTTCACAGCTGCACCAAGAAGAGTAACATAGTCTTTTATGAGGAGGAAGTGGCTGGCTGTACGCATGCGGGCGAACTGCTCCTCCAATGTAGATGTGACTTTGCCAATAGCTGTTTCCCATGTTGTTTCCACTTGACTCTCAGACAACAACCCGTCTGCAGTTCGAAAGACACGTTCTTCAACTATAAAAAACCCTGCCACCTGAGAAAGGAAGGGCTGATGGGACTCCAGGAATGGCTGTGAAGTGGAAATTTGCATGTCCAAGTTGAGTTGCATGAGCCTATTTTTGTAATAGTAATCTTTGAACTTCTCCTCAATACCAAGGCATATGTGCATGTGATGTGCCCGGTACACTGGTGTGAGATCAAAATCTAGTGCCGATTCCTCTTCTGTATTCTCCAGATCCAAGGTATACAGATGCTCATCTGGTCCGGCATGGCTATGCCCTTCTGCCTCTCTCTGTCGTGCACGCATTTCCTCCTCTCTTTGTCGAGCTAATGAGGCCTGGCTTATTGAGACCTGGCCAATCTGCTTCGCCATTCTTCTGATATGAACAAGCCAATCATTGAACTCGCTGGTAACTTTCTTCTCAATATGCAGCTTTATCAGTGGTATCTGCCTCGCCACCACCCTTTTAATAAGTTTCAATGGAATATTTTGCAAGTATCCCTTCTGAATCAGGTCCAAAGTCTTCAGTGCAGGATGAAACTTAGCTTCTGCAATATAATTGTTGCATGTCATGCACAAGCAGATGACCTTCACACATATCTTTAGAGTCGTTATGGCCTCCCCAACGTTTTTCTTGACAGAATATAACTCAAGAAGTTCATCAAGCTTCAGCAATTGAGCAGTGGAAGCCTGTTGTAGGTGAGAATTCTCACCAGATAGCATGCTCTTCAGCTCCTCAGCATCGACCAACACCCCACGTAGCTCATCAACAGCAACGATGAAATCCTCATAGTGAAGTCTACATAGTTCTTCAATTTCAACTTCCTTCTTCTTCACAATGCTTCTGAGGCAATGCGTGAGAGCTTCAGGTTTCCCAGACTCAAATGAGTGACGAATGATGGGACCCAAATCATCGCCATTTGCAATAAATGTTGCAAGGCTCGAACCAAGGCCTCCATCTCCACTATCCACTACACTCCTCTTCTTTCCTTGAGCAGGTTGAGCAGTCATTGTTCAACTGTAGAAAAGACGAATAGGTACACAAGTTTCAGAACTAAATGGCATTGTGGTTCACGTTTCATCCAAAATAAAACAAAGATTGGGACAACCATACAAGACCGGATGCATTATAATTTTAGACGAACAGCAGGAATAGCAATTAGCAGCTGATAAGCTTCATTTTAATACCAAGTTTGCCTAATGAACAGTTAGTATATTAGGCATTGGCACAAAGAGTTCAAAACGTGATGCTTCCTATATTCATTTCTAGACTTTAATTTACAATTGTCTACAGTAGTAATTATATTATTGACTAAAGGATTCATTGAAAATATTAAAAGCTGTACCTCAATACTTGTAGAAATTTAGAGACTCACACTTTACTATTCATTTTGAAAAGACTCGCGCATAGCAATTACCGCAATAACAGATCGACTAAAACAATGAATTGAAGAAAAAAACACTGAATAATCTGAAAGCATTTCATCTTGGGAAAGAGATTGATAATGCACAGTTCTGGAATGATTCTCACCACGTTACATATTTCCAGCAGATAAGGCCAAACGTAACAATTGATTCAACCGGACACAACGCGATAATCGCACAAGAACACAAAGGGGGCCAACTAAAGACCGAAGCACCAACAGCAATTGTAATAACATGTAAAGCACAAAACGGGGTATAGCCGTATAGAGAATGAAAAGCTTACAGTGCGAGGGGCTCGTCGCGGTGCCAACCCTGTCTAGCTGCAGGATCCTGGCGCCTGCCCTCGATGCAGGACGGTGACGAACTGCGAGACTGCAGCACGGAGACGCAGCGACGATGGAGGGAATGGATGGGCGCGCCTCGCCTGCCGCCCGAGCCCTCTCGCCGCCGCCGAGTCGGTCAGAACCAGAGGACCGGATACAAGCGAGCGCCGGGTTGTTTGCTCAGAGAATACAGCACCAGCGGACGCAATGGGATCTTGAGAAGGAGAACAGACGGACCACCAGAACGACGAGTTGATCGAAACAACAAACGGAAGCCGGAAGGGGGCAAGTAGGAGGAAGAGGAGAGAGAAAGAGAGTAGGAGCTCCAACGGTTGCAGGGGAGGGTGTCCATTTGCACGGGCGGGTGGATCCACGAAACTCGCGCGTTTACCGATTCTCCTCCGCCCGCCTCGCTCGGCCGCGAGGACAGGTGGTCGCCGCCGCCGGTGACGGAGGCGAGATCGACGCGGAGCAACGGCTCCATTTTTTATGGGCCACGAGGCGTTCTGTGGCCTGCCCGGACTCTTTCGGCCCATTCCGACGGCAGTGGTGAACGGGCAGCACGGGCGAAGAAGAGCCTAAAGTTCCCAAGTGACACATTCATTTCTCCTGCTTGAGTCACCTCATTTCACGTGGTTTGATTTTCTGGTGTTGTTCCTTTAAAATAAACGAATAGATCTTATCAAATTCACCCCATATACGGGTAACATCTGGCCCTACGGTCACGGATAACGAACATCTCATTCCATGTGGTTTGATTTTGTGCCTCACGCGGATGAAATTTCTACAAATCACCAACAAAATACTGCTCGTTACTAAGGTTGAACCCATCCTCTGATGGAGCGTAAACATGGTACTTATGAGGCCTTCCGTTTGCGAGACCTCGAGGTTATGATGCAGATGGTGCCGTGGCGAGCATGGCAATCTTATTATATCATCGTTAGGCAAGATGGCTGCAACATGTTTGTTCCCCATAATTACAAAAAAAAAAGGTTCAAAAAACAAAAGAGCTTCAGATTCTTGCGGGACCGTCGTCCTTTAGCTAACGGCGGATCGGATCGGATGGCAATAATGACATGGATGGAACTTAAGAGTCAACAATCCTATCATCTGCTGGATTTATTCCACGAAAAGGGCCCTCAATTCTTTTCCATCATTCCCTCCATCTGTCTCTGCTCAAGGCGGCACTTACTTATGCTTTCCTGCATGTAGAGCAGCGCATGA
