## Supplementary File 2 for "Population genomics uncover loci for trait improvement in the indigenous African cereal tef (*Eragrostis tef*)"

## >4A:14601886-14619069

CGTCGGGATCTGCCCTAGGGCTCATACGGTTTGCTGGTGTGGATCCTGGCCTAGGTTGATCCAGTGGAGGATCATCGTGGGTGTCGTGGTCCACCAACGACTCGTGGACATATAGGTTAGGGGCAATGGGATTTTCTCATATTTCGGTGGTGCTTCTGTTCAGGGTCTCGTATGAGTAGTTGGTGGTCACAAATCTAATCTCAGGAGATGATCGATGATGTTGGTCTTAAACGATTTCTGGAATCTCGTTTTGGTGTGGAATATTGGGGAATGGAACTAATTAAGTGATTTTTGTCAATTGTCTCAATGATTGGTTTATGTGATTGCGTCCCGCTTAAATCTTATGCTTAGTTGGTGTAAATTCTTCCCCAGCACACTGTAGTTGGTAATATAGTTGTGAAAAGAAGCTATGATTAGTACGTCAAACATGAATGCATCACTCGATCCTGGTGAAGTGTTATTGATATGTGTGCCCTAGACAGCAAAGAGGAAGGGGGCTTTTGTTGGTCTGAAGGCAATAAGGTGTTGAAGAAATTTCATCAAGGATCAAAGCTTGTGTCGCCTGTGGCATAGCCTTTTTGTTGAAGTTATTACATAGGAGGACTTGGGTAGTAGCAGCTATGTTGGTTGATGTCTTGCTGATCAGGCAGGCAGTTCTGAAGTTGTTGTGTTGCTGCAGGGGTAGGAAGGTGGCTGCGACTAATTTCTCCTGAGTGGTTGCAATGTGGTGTTGTTTGATTAATTTTTTCCTTTTTCATTTTGAAATGTCAGTCTTGTTTTAGTGGATAGAAGAATGAAGCCATTGCTTCCATGTTTCAACTTGAAAATTTCAATCTGTCTTATTCGTATGTATGATGTAAAACTGCTACATGACATGGAATGGCATAAGTGGTTACTTCTTTCATCGGTAAGGAAGCCTAGCTTGGTTCGACGAATAGTTATTGTTGTTCCATGGAATCTTAATTTGCGGCAAGTTCTCATATCTTAATCTCTTCTTTTTTTCTTCTTTTTCTTTTTGTGATGAGTACTCCAATCCACCTAATGTTGTCAGGAACTGCACTTTACAATTAGGTTGTATGTTCCTGAATAGTTTTGTTTTCATGGGGTAACATATTAGATTTCACATTCAATATAGATTGTGTTTTTAGTCATGACATCAATATCAGCATTAGTGATAATCTCTGCTACGATTATAATTACAAATGCAAATATATCCATTGCATCCTTCAATTCTACTGGTATAATTACAAAAAATAAAAAGGACTATGTAGTCATCATGACTGAAAAGTATAAAGTATGTAGATATCCTTGTTTTATTAGTGACAGTGATGATGTAGTGGAAAGATGTTAGTACTTGGTAGTTGGTACGATAGCATGACCATCCATCCTTAGGTTTATTTTCCTCAATAAATATTTGTGTATACAAATTTATGTTTGTTAGAAATGTCTGGACCTTATAACATGCGTATTTATATTCCCTGTCATCAGCAGTCCCCCGTCCTTTTAGTACTGCCTAGCACATGGTGATTGTACAATTATGGCTATGAGATAGGTTTGTGAATATCAACATGATTTTTATCTTTGCCATGCATTTTGATATTTCTAGTGTGCTTAGCATAGATATTGATTACGAATTTACTAAAATGCGTTGGGTGCATTTCTTGCTTTCTGTTCGAAACAATCAACTGCAGTAAGATCATCATTAAGATCCTGAATTTAGCTTTGTAAAATTCAGCTCATCGTGTCTGTTTACCAAGCTGTGATTCTGTATTGATTGCTAATGTACGATTTATTGGATGTTGTGCAGTGATGGAGAAGTGTCAAAACAACGCTAGGTTTGCATCCTTTAGTGACGCTCCGTTTGCTCTCTGCTGTAAGAAATCATGCATGGATTTTATTATAAGTACTATGTTCTTTGTCATTCTGGTTTAGTTTTGAGTTGCATCCATTAAGAGATAGTGGACACATAACTGGCATCCCTTCAAATGCATTAGCTCATTTAGGATGTGATGTAATATGGAGCTTGAAGTGGCAATCTGCTCTGCTGGCACAGATCTGAACTGAGCTGTTGACCAGTGAAGCAATACATTTTCCATTTTTCACAGCTCAAGTAAATGTTGATTGCAGTCCTTCGTAACTCTAGGAAACATTGATAGTGAATACTGATCCTTTCCACTATCAGACATTATTATATAAATGTTTGATTTTTGGAATCATATAGAAGTGCTTATCCATGCCTTGTGGCTTTAACCCTCAGTCCTCTTTTTGTTTTATCTGGATATAGGTATTATGTGTTGTACAGCTTCATGGGGAAACACCCATCTACTTTGAATTTTTTATTCTAAATTACCTTTTTTTGTTGTATGAGCTTTTGTCAATACAGCGGGACCTCTACTATATCTGTTCTAAGTTTGTTGCCTTCTTTTCAGGTGATCTTGGTAGCTCGATCTCGAATCTTGAGTAAGCTAAGGGCTACACATCCACAGTTGCTCTGCGACCAAGGACATCTCCATTAGGAAGCTGGCCTCCATGAAGGTCAAGATCTTGCAGTGGCACGCGGTGGCTTCATGGACGTGGGATGCGCAAGATGAAACATGTGGCATATGCAGGATGGCATTTGACGGCTGCTGCACTGACTGTAAGTTCCCTGGTGATGATTGCCCGCTCATGTGGGGTGCGTGCAATCATGCTTACCATCTCCACTGCATACTGAAGTGGGTCAATTCGCAAACATCAACGCCCCTTTGCCCCATGTGCCGCAGGGAATGGCAGTTTAAGGGCTAATGCACATTAGTGGCGTATCTGGAATGTATCAGCAGGGAATGGCAGTTTAAGGGCTAATGCACATTAGTTTAAAGCTAGAATGTATTGTTGGACTGGAATTTTAGCGATGCTGTCACTTTATTGTGCTAAAAAGCTACAATCCTCATTGCTTATTAGCATCTGCTTAAGTGAACTTGTGCCGACTTGAAGTTGTGGCCAAGCAAATGAAATGATAATATTTCAGAGGCGTTAAGCTTTCTGACCTCATTGCTATTTTTTGTGGTCCTATTGTTTACTTTCTGAAGGATTTCAAGTGTCAACTTTTGGATTTTATTCTGGTAAAGTAGTACTAACAAGCATGAAGCTGTATATGGCTTTGCTTGCTTTTATCTCTGTTCTCTTCTTCCACAAAAATAGTGATCCATAGGTTATGAGGTGTCACTTTAAGCTCTGCGCGCAATTTCAGTAGGCATGTGTGTATTATTTAGTCGTATACTAAAAAATAAGTATCTAAAATTTGTTTAAAGAAGGAACGAACGGGACATGCAAGTTTATGCTATTGTGGTAAATTATATCCACCGTTCCTTCCAAAACTTTGTCCCTTGCAATTTGTTTCTTCCGTGCCAATGTAGTAATGGCAATATGCCAATAAGTTAAACCTTATTAGCTTGTGCAGAAGCATGGATTGACTGTTAGTGCTTATCTAGCTTCAATCTGGACAGATTGGAATGTGCATTGAGTTACTTTCCCTATTAATAGGGGGGGGGGGGGGGGGGGGGGGGGTCATATGAAAATTGGATGTATAGCAGTTACGCGTTCATAGACTTGTTTTGTTACTTGTCCAAAAATCACTCAAGGAATGAAAAGTTGCACTTGGTGTTTCCCCCCCCCCCCCCCCCCCCCATCTGCTTGAGATATATGGAGCCACCGCAAATATTTGTTTGTGACCAAACAAGGGCTATGCCAAAACTGTAAATAGTCAAACAATATCCTTCAAATTTTAAGATCTTGTTTAGCATGCAATTGTAAGTTGGATTACCATTTAGGATATATTGACAGTTAAATAATTGTTAACCTTTGTTTGTTGGACGGACCCCATGCAGCCTCAGGCTTAACCTTTGTTTTTAGGTTCCGTTGGCAACCAAACTAGCCAAATCGCTGGCGAGAAGTTTGGCAAGGTTGAGATTATGAGATGGGCAGTAAACCATACACGCGCTTAGATCTGGGGGTAACGTAGGTGTGCTAACAGTACATTCTAACCTGAGTCACAGCGCAACATTTACAAGGAACCAACATTTTTCTTCCTAGCCAAACGATGGAACTCACCGATTTAATGGGTCCAGGAAGGAGCACGCCTGGTACCTATCAAGAGAACTTGCTACCGCCTGCAAATACAAAGCCCGTCTAGCTCAGTCGGTAGAGCGCAAGGCTCTTAACCTTGTGGTCGTGGGTTCGAGCCCCACGGTGGGCGATATCTGTTTCAATTTTTTTTTTTGAACCCGTATCTTTTCTTCTTCTTCTTCTTATGTGATTACTTCAAAATATACGGTGGAATTCTGGAATGTCACATCCTTTTGTTTCTTCTTATGATAGTGCAAATACGCACGGTGGACGATATCTGTTTCAATTTTTTTTTTTTTGGAACCCGTATCTTTTCTTCTTCTTCTTATGTGATTACTTTAAAATATACAGTGGAATTCTGGAATGTCACATCCTTTTGTTTCTTCTTATGATAGTGCAAATACGCTGGAATTCTGCCTTTCGACATGTCCGATTTGCTTCTTTATTCAGATCATGCTGATCCTCCTCGGTGAGCTTAATTGGCATCATGTATAGCCAGCCATATCTAATCTTGTGTACAGTTGATGATGATGAGCTTTGCCCTTAATCCTCACTAACTGTCACCACGTCGCGGTACAACCTAGAGCGACATCAAAACCATGCAAACTCCACGTCGTCTCCTGCCGGCTCCAACGTCAGCAGCGCCTCCATGTCACCCTCGCCGCCGACGAACTGGCCGGCCGGCCCCACCACGCCGTCGCCGGCCTCCCCGTGCCACGGGCTCAAGAAACCCAGCTCGTCCATGTCGAAGTCAAGGTCGATGGAGATGTCATCCTCCGGCAAATGCTCCGGTCGCACCTCCGCCACCGCCGGCGCCTCCGGTGGACGGCCGCGGCTCACGGCCGGCGAGGCGGCGGCGCGCGGCAACCTGGCGGTGCACCGCATCGCCTTGGTCCGGACCGGCGGGCTCGTGTTGCCGCCGTCCGGCGACGGAGGCGCGGCGTCGCTGCTGGTTCTGGCCGGCTCCGAGGACGGCCGGCTGCGTACGACGACGGCCGACGCTGCTGCGCTGGCTTGATGATGACCCTGTTTCGAGTGGCCGCCCATGACCTTCTTGCCCAGCGTCGTGTTCCAGTAGTTCTTGA***TGTTGGAGTGTGTTGGCCCATATAGAGGCCCATGTATGTTGGATATATATATACCCACCCATCTAGGGTTGGAGGAATCAAGTCGTCCAGTACCCTAACTCTAACA***TGGTATCAGAGCCTCTCTCTCTCCCTACCCAACCCTAGCCGCCGCCGCCGCCATGGCCGGCTGGCCGCCGTCGGGATCACCGCCCCCCGTCGTTGCCGCCGCAGGTGCGTGCGCTGCCGCAGGCCACCCGCCGCCGCCCCCCGCCGTCGCCGCGGGTGCGGGCGCTGCCACGGGCATGGACCTCGCGCCGCCGCCCCCCGCCGCCGCTGCAGGTGACCCACCAGCCGATCCTCGTCTCCGACTCGATGCCAGCGATCCGCTGGTGGCCCAGCTCCACCTCCAGGCAGTCGGTGTCCAGAACGTCCGCGCTATGGTCTCCATCGTCCTCGACGTGTCGTCCACCAACTACGGCCGTTGGCGCGATCAGATCCTCCTCGCCCTTCGCCGCTACGCGCTCGACGACCACGTCCTCACCGACACTCCTGCTCACGCGCGCGGTGGTTCGTGGATCTACCTCGACAGCGTCGTCGTGTCCTGGATCTTCGGGACCCTCTCCCCAGATCTTCAGGAACTCATCCGGGCTTCTGAGGACACCGCACGCCAGGCCTGGCGGGCTCTCGAGAACCAGTTCCTCGACAACGCCCAGGCCCGCGTCCTACACCTCGACACCCAGCTCCGCACCTTCTCCCAGGGCGACCTTTCTGTGACCGACTACTGCAGGAAGATGAAGGGCATGGCGGATGGGCTTCGGGACCTCGGTTCTCCTGTGGACGAGCAGCTTCTGGTCATGTACCTATTGAGGGGTCTCAACAGCAACTATGACCTCCTCAAGACCTGGCTACCTCGACAGCGCCCCTTCCCCTCCTTCCAGGACGTGAAGAATGAGCTCCGCTTCGACGAGATCACCCGGGGCCTTGCACCAGGACCTGGATCCTCCACCACCGCCCTCGTTGCTGCCCCGCCCACCACTCCACGGGTTGCCGCTCCTCCCCCTGGGCCGAGCGGGGACGGGGGTGACCATCGTCGTCGCCGGAGACGTGGGGGCGGCCGCGGGACCTCTACACCGACTCCTACTCCGGCGGGTCCGTCCTGGCCATCCTTCACCAACCCATGGTCGGGGCGCATCTCCATGTGGCCGTTCCAGCCCGCCGGAGCCGGGGCTCGTCCTCCGCACCGGCCCGCTGCTATGCTCGCCGGCCCTGCTCCCCCGCTGCCCTGGGCTCCTCCCGCTCCGGTCGGCCAACCTCCGACCTGGCCGGGGGGGTGGACTCCTCCTGCTCCGACTTGGCCGGGGGGGTGGCCTCCTCCTGCTCCGATTTGGCCGGGGGGGCAGGACCCGGCGGCGTTGGCCCAGTCTCTCAGCACGATGCCGCTGACCCCACCAGCCGACTGGATCGCGGACTCGGGAGCCTCGTACCACACCACGCCCGATGCCGGTATACTTTCTTCTGTCCGACCTCCCCACCCTTCCTGTCCCTCTGCCATTATGGTGGGCAATGGTGCCTGCCTTCCTGTCACCTCTGTTGGCACTGTGCCGGGTTCTTTTCCCCTTTCTGATGTTCTTGTTGCTCCCCAGATGGTTCACAACCTCCTTTCCATTCGCAAATTTACTACCGACAATTCTTGTTCTGTTGAGTTTGATCCTTGTGGTGTTTCTGTGAAGGATTTGGCTTCCCAGCGTCCACTACTTCGATGTGACAGCACGGGACCCCTTTACACGATCCGCCTTCCCGCTTCCGCTCCTTCATCGCCATCTTCTTCACCATTGGCCTTTGCTGCTACATCTTCTTCTACCACCTGGCATCGCCGGCTTGGACATCCCGGCCGTGACGTTTTGGCCCGACTCTGCCGCAGTGTGGATCTCTCTTGTACTCGGGCAGATGATGAGCACCTCTGTCATGCGTGCCAATTAGGTCGTCATGTTAGACTTCCTTTTCCTACTTCCCAGTCGCATGCGACTCATGTTTTTGATCTTATTCACTGTGATCTGTGGACTTCACCTGTAGTCAGTATTTCTGGGTATAAGTACTACTTGGTGATTGTTGATGACTTCTCTCACTATTCTTGGACTTTTCCTTTACGCGCGAAGTCTGACACTTTTCCCACCCTCCTCAACTTCTTTGCTTGGGTCTCCACTCAGTTTGGCCTCACCGTTAAAGCTGTTCAGTGTGACAACGGCCGTGAGTTCGACAACCACACCTCCCGAGCCTTCTTCCTCTCCCGCGGCGTTCACTTGCGCATGTCTTGTCCGTACACATCTCCTCAGAACGGCAAGGCAGAGCGCATGATCCGCACTACGAATGATATTGTTCGTACCCTACTTGTACAGGCCTCCCTCCCGGCCCGCTTCTGGGCCGAGAGCCTACACACCGCCACCTATCTCCTCAACCGTCTCCCTTCCACTGCCTCCCCTGCTCCCACTCCCCACCACGCCCTTTTCGGCTCCCCTCCTAGATATGACCACCTCCGTGTCTTCGGGTGCGCCTGCTACCCCAACACCTCTGCCACTGCTCCTCACAAGTTGGCGCCCCGTTCGACCCGGTGTGTATTTCTCGGGTACTCTCCGGAACACAAGGGGTATCGCTGCTATGACCTTACATCTCGTCGGGTCCTAATCTCTCGGCACGTCGTGTTTGATGAAACTGACTTTCCCTACTCCACCTCCACCACTCCCTCTCTGGATCCCGAGATCGGGTCTGTGTTTCCGACTGACCCGGTGGCCCAGTCACCTGTCTCTGTGTATCCTTTACCTACAGGTCTTCAGGGCACCCCGACCCCGAGTCGGCCGCGCTCGCGATCCCCGCCTCGCCCGACCCCGAGTCGGCCAGGTCGGGCGGACCCCACCCCGAGTCGGCCGCCCTCGCGATCCCCGCCTCGCCCGACCCCGAGTCGGCCAGGTCGGGCGGACCCCACCCCGAGTCGGCCGCGCTCGCGATCCCCGCCTCGCCCGACCCCGAGTCTGCCGGGTCGGGCGGACCCCACCCCGAGTCGACCGCGCTCGCGATCCCCGCCTCGCCCGACCCCGAGTCTGCCGGGCCGGGCGGACCCCACCCCGAGTCGGCCGTGCTCGCGATCTCCGCCTCGCCCGACTCCGCCTCGCCCGACCCCGAGTCGGGCGGGCTCCCGACCGCCCCCGAGTCGACCGGGCTCGCGATCCCCACCTCGTCCGACCCCGACACCTTCTCCCGCGCGCTACGCCCGGCCTCCGCTCGTCTACCAGCGCCGGGCGGCACCAACTCCACCGGTGCCTCCGCCCCCACAGCCGACTGCGGCCCCGTCTCGGTCCCTCCCCCGCCTCAACCCGGTGGTGTACCACCCGCCTGTCATCCATCGGGATCCCGGACACACTCATCCCATGGTGACCCGGCGGGTGGCGGGGGTTCTTCGACCTAAGGTCCTCACTGCTACTGCGGACGAGCCGGCGGTCTCGCCGATTCCTACCTCTGTCCGTGGCGCCCTGGCTGATCCTCACTGGCGCCGGGCGATGGAGGAGGAGTATGCGGCCTTACTTGCCAACCAGACGTGGGACCTCGTTCCTCGTCCACCTGGTGGCAATGTGGTCACAGGGAAGTGGATCTGGACTCACAAGCGTCGTGCTGATGGCACACTAGATCGGTACAAGGCTCGCTGGGTTCTCAGGGGATTCACCCAGCGGCCTGGAGTTGATTATGATGAGACCTTCAGTCCCGTGGTGAAGCCGGCTACAGTCAGGTCAGTTCTGTCGCTGGCTCTCTCTCGCTCTTGGCCTGTGCATCAGCTGGATGTGAAGAATGCTTTCCTGCACGGCACTCTCTCTGAGACAGTTTACTGCAGTCAGCCAGCGGGGTTTGTGGATCCTGGTCGTCCTGACATGGTCTGCCGACTCAACAAGTCCCTCTATGGTCTGAAGCAGGCACCTCGGGCTTGGTACTCTCGGTTTGCCTCATTCCTGGTCTCTTTGGGTTTCACTGAGTGCAAGTCGGATACCTCGCTGTTCGTCTACGGCCATGGCGCTGACACTGCATATCTACTGCTGTATGTTGATGACATCATCCTCACAGCGTCCAGCCCGGCCTTGCTCCAGCGGATCATTCAGGCCTTGCAGCGCGAGTTTGCTATGAAGGATCTGGGTCTCCTTCATCACTTCCTGGGGGTCACTGTTGAGCACCGTCCTTCTGGCCTGTTCCTTCATCAGCGGCAGTATGCTCTTGATATTCTGGAGCGTGCCGGGATGTCTGACTGCAAGCCGAGTTCCACCCCCGTTGACACTCAGGCCAAGCTATCTGCTGATTTGGGCGCTCCGGTTGCTGACCCGACTGCCTACAGGAGCCTCGCTGGGGCTTTGCAGTATCTCACCTTCACCAGGCCTGACATCGCTTATGCCATCCAGCAGGTCTGTCTTCACATGCACGACCCCCGCGAGTCCCACCTGGCTGCTCTGAAGCGCCTCCTCCGCTATGTTCGTGGCACAGTTGACTATGGCCTGACTCTTCACCGGTCTCCTTCTGCTGAGCTGGTTGTCTACACGGATGCCGACTGGGCTGGCTGTCCGGACACTCGGCGTTCCACCTCGGGCTACGCAGTCTTCCTCGGCGGCAACCTTGTCTCGTGGTCGTCCAAGCGACAGCCCGTGGTCTCCCGCTCCAGTGCCGAGGCGGAGTACCGGGCTGTGGCCAACGGCGTCGCTGAGGCGACCTGGCTTCGACAGCTCTTGGCGGAGCTCCACACTCCACTCGCCAAGGCTACACTTGTCTACTGCGACAACGTCAGCGCGGTGTACCTCTCCACCAACCCGGTTCAGCACCAGCGGACGAAGCATGTCGAGATCGATCTCCACTTCGTTCGGGACTTCGTCGCCATTGGGCAAGTGCGGGTCCTCCACGTTCCCACCACCTCCCAGTTTGCGGACATCTTCACCAAGGGTCTCCCCTCCTCGATCTTCTCGGAGTTTCGATCCAGTCTCAACATATCCGGTGGCTAGTTGTGACTGCGGGGGGGGGGGGGGGGGTGTGAGTTGTTCTTTCTTGTGTCCAGTCCTGAACACCGCTGCGACGGTTAGTTCAGACTGCGGGGGGG***TGTTGGAGTGTGTTGGCCCATATAGAGGCCCATGTATGTTGGATATATATATACCCACCCATCTAGGGTTGGAGGAATCAAGTCGTCCAGTACCCTAACTCTAACA***CTTGATCTCATTGTCCGTTCGCCCCGGCAGCCGCCCGGCGATCAGGGACCACCTGTCCGGCGATGTCGACAAAATGCAAACACGTACGAAATCGTTTGTTAGTCACTGCTGGATTGCAGTTTCAGTCATAGCATTAGATGTGTCAATCATCGATGGTGCCAGCCTGCTGTTGTTACTCGATCTGTCCCGTTGATCACATCAAGTTGTTTTTGAGGAAACTAATCACATCAAGTTATGAGCAAGTGGTAGTAGAAATGAAATACTTGCATTGGCAGCACAGTTAAAGCTTTCCATTTCTGGAAGGACAATATCTTTTGCGGTCCGTGAAAAGTTTTGGAATTTTCAGGGTGAAGTAAACGCCAGTTAGGAAGTTGTTACCCGCGCCAGTGGCCTTTCTTGAGTGCGAGATATTCTCGTGTTTCTGTTATATATTTAGGGAGGGAGGGACAGGGCTACTAGCTGTCATCATCAACTTTGCTGACCTAATAACTGGTTCATCTTCAGAGGGTAGGAGGCACTGCACTGCCAACGAATACAAGATCGATCTTCTAGAGCGAGAGAGCTAAACAGGAAAAAAGAGGTGCTGACTGATATGGACAAAGTGGCAGTAGTGGTTATGATAACTTGAGGCAATTGGCCGTTACGCTTGCCTGCTGGCCTACTTTGTCTGCTCAGATCTCTACCACAGCAGGACAAGGAAACGTTTCCAGCAGAAAGAGTTTAACATGCAAGCTAATGTGTTGTTGTATGCATGAATAACGCAATTACCCTTCTGTCTTCTATACCATACAAACTAACATTTCTTTGAACAATAACAAACCAAAATTTAGTATCTTTAGGATTTAATTTGCGAAATGCAAATTGATGGAAATGGTCGAAATATGGATAACCACATACCTGTTGCCGAGGAGCCTGTGCAGCCTGATGATGAGCTCCTCTTCGTCGTCGGAGATGTTGCCCCGCTTGATGCCCGGCCTAAGGTAGTTGAGCCACCGCAGCCGGCAGCTCTTTCCGCAGCGCTTCAGCCCTGCAACACCTCGATGTCGTCAGTGTTGCGCATTCCCAACAGTCAGCGAATTACTGTACTGAATTTGTCATGTCACATTGCATTTCGTCCTTGCATCGCCCTACTTGTAACCGATCCAAGCGTAGTTGCTCTCCGTTCCAGTTACAGAGGTCTATGTAGACTCATAAGGTAATCTAATCAGCATTGAGTATGCCTTTGATTAACAGTTTGTTTAGTGAATACAGTGAAAAATAGACCTGTTGTCATTCGATCCAAGCGTACTTGCTCTCCGTTCCAGTTACAGAGGTCTATGTAGACTCATAAGGTAATCTAATCAGCATTGAGTATGCCTTTGATTAACAGTTTGTTTAGTGAATACAGTGAAAAATAGACCTGTTGTCATTTCATTTTCATCCTAATATCTGATCTAAATGTAACTTAGCATGCATGAACTGCAAGACGTGACAAATCCATACATAGACGATGATAAAAAAATCAGAAATGACTAAAAGAGAAGATAAGAAACAGGGGACGGGGTAAGAATTTCAGGATCAGACCAACTTCCCGCATTGTGAATAAATTGATGTTGATTGTCGATGAAGAGAAATCGGCAAGAGAGAGGGAGGAGGTGTCACTGACCAGCTCTTTTGGGGAGGCATCCCCACTTGCCCTCTCCATGCTTCCTGATGTAGGAGACGAGGATGTCATCCTCCATTGCTGTCCAAGCTCCCCTGTTCAGCCCCTCCTTGAGGCAGCATGGTTTCCTCCCCATCTCTCTCTCTCTCTCTCTCTCTCTCTCTCTCTCTCTCTCTGTCTCGGTTAGATGGAGGGTTATGCTCTGTGATTGCCACTGAGGCCCCTATTAAACCAGGTGCAGAGCGGTCCGTATTCCATGCCCATTTTTTTTTAAAAAAAAAGAGATAAATATTTGTGATATAGCCCCTCGAATGGACATTTATGCCCTACGGGCACTCTGTTTTCGCCCACACAGCTACTTCTGATGAGACGCTCGTTATGAAGGCAGTTTCTAATCAGAAATCAGGAAGCTTTCGCTTGTTTATGCACATACAGGGAATGCTTAAAATTTTGACCAGCATAAAATTTCAGAAAATGATTTACAAAGCACCGAATTACTTAAGACCATATCGAATACGACAAGTGCATGTTCTAGATGCAGATGCCAATGTGGCACCTGCAACAGGCTTGAGAAGTACCGAGAGGGAAAACATATTGACACACAATTTTTGTCAACTCTGATTCTTCTTCGTTTATCAAGGTGTCAGTGGCCATGGTAATGAAAAGAAAAATCTGATCTACAACCATGGTATAGGAAACTACATCCTATCTGGGAATATACTCCTACTAATTCACAAATAGTCAACACTTACCTATTAATATAAATCTAAAAGGTGGATAATAGCACAGAATTTATTTGCAAATGCAACAAGCATAACCTTAGCTAAAATCCTTAAGCCTTCTCTTCAAAACATCCATGGACCTCTTGCGAGCACTCTGCTTCGCGTTACGGTTCGAAAGGCTCCCAAACAGTCCGTCAGCAGCGTCCTTGTACTTGTCACAAATGATGGAAAGCTTCTTGTAGTCCAGTGATCCGTAGTTCCTCTGTCTTATCACCGGATTCATGAAGTTCTCTGGTTGGCTACTGAGCAGGAGGTTGACTAGCTGCCTTATTTCAACCAAACAATCTCTGAAAGTTGTTTCCTTTCCCGAAACCGACAACCCCATGCTTTCGAATTTCTCATCTGCAAATGCCTCCAGGATTTTCAAGTCCATATCAAGACCTAGAACTGCATTAGCAGTGAACCTTTTCACACCTTCGTTAAGAAGTGTTGTCATTATCGAATCTGAGATATGGCTCATTATACCAGAGACCACCTTGTATAGAGCTTCCAAGGGCAAAATTTCCTGTGCTGTTGATACTTGAGTTTCGAGATAGATTAGGACCTCATTCATATAGTCGTTTGAAATGTCTGGAGCTTCTTCTGCTATCCAGTTGACGTTTTCCAAGAGTATCATAAACTCATCGACCTTGAAATTGGCCAGGTTTATCAGAGCATTGTATACTGCATTCTGCGAGGCTTTCAAGACAGCTCGAGCAGTTAAACTAGAATGAGACCTCTCCGCAATGCGTTTTGGTATGCCACACAGATGAGCAGCATGCAAGAGGAACATGCCACATGCTTGCTCAAGAACAGAAATGTTACCAGCGAGCTGCATCATCTGTGACATGGCCAATGAGCGAGCATAGATCATGTTCAGAAGGCTGTCATTTAAGACCTCAATCAACAGCCTGTCTAGGTATGCTTTCACCACATCATACATGTTCATGAAGCCACCATATGACAAGTAACTGACGGAATCTTCAATGAAAGACCGAACGATGCGACAAATTTCTGGAACAGAAGAGGAGAACTGCGCCACATATGGGAAATCAGGGGCTGCATCATCAGTTTCCAAACAAAATGCTGTCACATTCATGTTATACTCGTATTCCTTCTTTATAACCATCTGCTCATATGAGTCATTAATAAGGATGTCATCCATCTGCTTTCGGCATTCAAGAAGAAGCAACTGGTGGTACTTGTCCCTGCTTTTCTCAAGAACTTCAATCAACTGTGTAATTTGATAACCATATTTCTTCACAGCTGCACCAAGAAGAGTAACATAGTCTTTTATGAGGAGGAAGTGGCTGGCTGTACGCATGCCGGCGAACTGATCCTCCAATGTAGATGTGACTTTGCCAATAGCTGTTTCCCATGTTGTTTCCACTTGACTCTCAGACAACAACCTGTCTGCAGTTCGAAAGACACGTTCTTCAACTATAAAAAACCCTGCTACCTGAGAAAGGAAGGGCTGATGGGACTCCAGGAATGGCTGCGAAGTGGAAATTTGCATGTCCAAGTTGAGTTGCATGAGCCTATTTTTGTAATAGTAATCTTTGAACTTTTCCTCAATACCAAGGCATATGTGCATGTGATGTGCCCGGTACACTGGTGTGAGATCAAAATCTAGTGCCGATTCCTCTTCTGTATTCTCCAGATCCAAGGTATACAGATGCTCATCTGGTCCGGCATGGCTATGCCCTTCTGCCTCTCTCTGTCGTGCACGCATTTCCTCCTCTCTTTGTCGAGCTAATGAGGCCTGGCTTATTGAGACCTGGCCAATCTGCTTCGCCATTCTTCTGATATGAACAAGCCAATCATTGAACTCGCTGGTAACTTTCTTCTCAATATGCAGCTTTATCAGTGGTATCTGCCTCGCCACCACCCTTTTAATAAGTTTCAATGGAATATTTTGCAAGTATCCCTTCTGAATCAGGTCCAAAGTCTTCAGTGCAGGATGAAACTTAGCTTCTGCAATATAATTGTTGCATGTCATGCACAAGCTGATGACCTTCACACATATCTTTAGAGTCGTTATGGCCTCCCCAACGTTTTTCTTGACAGAATATAACTCAAGAAGTTCATCAAGCTTCAGCAATTGAGCAGTGGAAGCCCCTTGTAGGTGAGAATTCTCACCAGATAGCATGCTCTTCAGCTCCTCAGCATCGACCAACACCCCACGTAGCTCGTCAACAGCAACGATGAAATCCTCATAGTGAAGTCTACATAGTTCTTCAATTTCAACTTCCTTCTTCTTCACAATGCTTCTGAGGCAATGCGTGAGAGCTTCAGGTTTCCCAGACTCAAATGAGTGACGGATGATGGGACCCAAATCATCACCATTTGCAATAAACGTTGCAAGGCTCGAACCAAGGCCTCCATCTCCACTTTCCACAACAGTCCTCTTCTTTGGTTGAGCAGGTTGAGCAGTCATTGTTCAACTGTGGAAAAGACAAATAGGTACAAAAGTTTCAATGCATATGTAAAAGCACAGAACTAAATGGCATTGTGGTTCACGTTTCAGCCAAAATGAAACAAAGCTTGGGACAACCATACAAGACCGGATGCACTATAATTTTAGACGGACAGCAGGAATAGCAATTAGCAGCTGATAAGCTTCATTTTTATTACCATTTTTGCCTAATTAACAGTTAGCATATTACGCATTGGCACAAAGAGTTCAAAACGTGATGCTTCCTATATTCATTTCTAGACTTTAATTTACAATTTCCTACAGTAGTATTTATATTATTGAGTAAAGGATTCATTGAAAATATCAAAAGCTGTATCCCAATACTTGTAGAAATTTAGAGAGTTGCACTTTACTATTCATTTTGAAAAGACTCGCGCACAGCTACTACCGCAATGAAGCTAGACTAAAACAATGAATGAAAAAAAGAAACACTGAATAACCTGAAAACATTTCATCTTGGGGAAGACATTGATAATGCAGATTTCTGGAATGATTCTCACCACGTTACGCATTTCCAGCAGATCAGGCCAAACGTAACAATTGATTCAACCGGACACAATGCGATAGTCGCACAAGAACACAAAGGGGGGCAACTAAAGACCGACGCACCAACCACAATTGTAATAACATGTAAAGCACAAAACGGGGTGTAGAGAACGAAAAGCATACAGTGCGAGGGACGCCGCGGGGCCAACCCTGTCTTGCTGCAGGATCCTGGCGCCTGCCCTCGATGCAGGACGGCGACGAGCTAAGAGTACCGCGAGATTGCAGCACGAAGACGCAGCGACGATGGGCGCGCCTCGCCTGCCGCCCGAGCCCTCTCGCCGCCGCCGAGTGGGTCAGAGCCAGAGGACTGGATACAAGCGAGCGCAATGGGATCTTCAGAAGGAGAACAGACGGACCACCAGAACGGCGAGTTGATCGAAACAACAAACGGAAGCCGGAAGGGGTCCCGCAAGAAGATGGAGCCCGGGGAACCAAAGAACTGGGGGGAGGAGGAGAGAGAAAGAGAGTAGGAGCTCCAACGGTTTGCAGGGGAGGGCGTCCATTTGCACGGGCGGGTGGATCCACGAAACTCGCGCGCTTACCGATTCTCCTCCGCCCGCCTCGCTCGGCCGCGAGGACAGGTGGTCGCCGCCGCCGGTGACGGAGGCGAGATCGACGCGGAGCAACGGCTCCATTTTTTATGGGCCACGAGGCGTTCTCTGGCCTGCCCGGCCTCTTTCGGCCCATTTCGACGGCAGGGATGAACGGCAGCACGGGCGAAGAAGAAAATCCCCAAGTGACACATTCATTTCACGTGTTTTTTTTTTAGACCAATTCATTCACGTGGTTTGATTTTCTGGTGTTGTTCCTCTAAAATAAACGCACAGATCTTATCAAATTCATCCCATATATCGGCAACATACCGCCCTATAAAGAACATCTCATTCCATGTGGTTTGATTGTGTGCCTCACACAGATGAAATTTCAACAAATCACCAACAAATTACTGTTCGTTAAGGTTGAAACCATCCTCTGATGGAGCGTAAAACGGTAAACATGGTACTATGAGACCTGTCGTTTGAGAGAACTCTAGGTATGATGCAGATGGTGCCGTGATGAGCATGGCAATCTTATTATATCATCGTTAGGCAAGATGGCTGCACCATGTTTTGTTCCCCATAATAACAAAAGAAAAAGTTCAAAAAACAAAAGAGCTTCAGATTCTTGCGGGACCGTCGTCCTTGAGCAACGGCCGATCGGATTGGATGCCAATAATGACATGGATGGAACTTAAGAGTCAACAATCCTATCACCTGCTGGATTTATTCCACGAAAAGGGCCCTCAATTCTTTTCCATCATTCCCTCCAACTGTCTCTGTTCAAGGCTGCACTTATGCTTTCCTGCATGTAGAGCAGCGCATGA
